# The ‘false transience’ hypothesis: Do we need to reconsider the seed persistence classifications of soil seed bank studies?

**DOI:** 10.64898/2026.10.16.759570

**Authors:** Judit Sonkoly, Viktória Törő-Szijgyártó, Eszter Ruprecht, Katalin Tóth, Gergely Kovacsics-Vári, Szilvia Madar, Edina Tóth, Péter Török

## Abstract

**Background and Aims:** Seed bank persistence data is typically indirect, inferred from the presence and distribution of seeds in the soil. However, this approach frequently underestimates seed longevity, because dormant and low-abundance seeds can remain undetected. Here, we introduce and test the ’false transience’ hypothesis which proposes that a subset of species classified as having transient seeds based on soil seed bank (SSB) studies actually possess the capacity for long-term persistence. Their perceived transience stems from methodological limitations and ecological filters, rather than physiological constraints. We also aimed to assess how the seed mass and seed production of ‘false transient’ species compare to the full range of tested species.

**Methods:** We focused on species considered to have transient seeds based on SSB studies. We conducted a germination experiment with 103 species using dry-stored seeds aged 10-29 years. We then assessed how the seed mass and seed number of germinating species relate to the full range of tested species.

**Key Results:** From the 103 tested species, eight species (∼8%) germinated. The majority of species germinating after >10 years had large seeds and low seed numbers compared to the full range of species, and all germinating species had seed numbers below the median of the tested species.

**Conclusions:** Our results provide partial support for the proposed ‘false transience’ hypothesis, showing that nearly 10% of supposedly transient species can persist after more than a decade of *ex situ* storage. Our results also suggest that ’false transients’ are frequently large-seeded with low seed production, therefore being more likely to be missed in SSB studies and erroneously classified as transient. We can conclude that relying solely on field SSB data to infer seed persistence can be misleading, suggesting that the widely observed negative relationship between seed size and persistence may also require reconsideration.

## Introduction

Persistent seed banks play a pivotal role not only in community resilience (Kiss *et al*., 2018), but also in providing ecological memory (Sun *et al*., 2013), and in the long-term maintenance of biodiversity (Royo and Ristau, 2013). However, compared to aboveground vegetation, we still have disproportionately little information on the dynamics and functioning of the soil seed bank (SSB). Although the formation of a persistent SSB is a complex process driven by multiple interacting factors (Long *et al*., 2015; Jaganathan *et al*., 2019), high potential seed longevity is undoubtedly central in this respect. Knowledge of seed longevity, i.e., the time period during which seeds of a plant species can remain viable (Saatkamp *et al*., 2009), is critical for understanding how population dynamics respond to environmental fluctuations (Tognela *et al*., 2025). Seed longevity was hypothesised to have evolved as an adaptation to climatic variability, particularly arid environmental conditions (Probert *et al*., 2009). Thus, extended seed longevity may also be a key trait for plants in the face of current climate change. Some methods can directly measure seed longevity, most importantly (i) carbon dating of viable seeds (Moriuchi *et al*., 2000), (ii) experimental seed burial (Telewski and Zeevaart, 2002; Pakeman *et al*., 2012), and (iii) viability testing of dry-stored seeds of a known age (such as seeds from herbaria and seed collections, see e.g., Milberg 1994; Molnár V. *et al*., 2015). While these direct methods can determine the potential longevity of seeds relatively accurately, seed longevity data assessed by these methods is available only for a very limited number of species.

In contrast to direct measures of seed longevity, field data on SSB persistence is available for a relatively high number of species (see e.g., Thompson *et al*., 1997; Wang *et al*., 2024). However, SSB persistence data is generally indirect, typically inferred from the presence, absence, and the vertical distribution of seeds in the soil (Saatkamp *et al*., 2009). In such studies, a species is generally categorised to have transient seeds if it is present in the aboveground vegetation and its seeds are either not detected in the soil or they are present only in the upper layer of the soil and absent from deeper layers (Thompson *et al*., 1997). However, this approach frequently underestimates species’ potential seed longevity (Thompson *et al*., 1997). Realised (ecological) seed persistence is often cut short by environmental pressures such as germination cues, fungal seed pathogens, or seed predation (Chambers and MacMahon, 1994). Furthermore, even realised seed persistence may go undetected due to low sample volumes or seed dormancy, which can confound germination-based SSB assessments (Saatkamp *et al*., 2009). Methodological limitations also stem from physical and demographic constraints. The size and shape of seeds strongly influences their ability to move downwards in the soil (Bekker *et al*., 1998). Furthermore, species that have low abundance in the aboveground vegetation or have low seed production are frequently absent from soil samples, leading us to mistake them for transient species, because of their failure to reach deeper soil layers or attain detectable seed quantities (Thompson *et al*., 2003; Saatkamp *et al*., 2009). On the one hand, this discrepancy highlights the fundamental gap between realised SSB persistence and intrinsic physiological potential, i.e., the maximum potential longevity determined by internal biochemistry and seed coat properties, isolated from ecological pressures. On the other hand, it also sheds light on methodological limitations.

Based on these assumptions, we propose the ’false transience’ hypothesis, which proposes that a subset of species classified as having transient seeds by field SSB studies actually possess the physiological capacity for long-term persistence; their perceived transience under field conditions is caused by methodological limitations and ecological filters, rather than physiological constraints. It implies that several ‘false transient’ species may, in fact, be represented by persistent seeds in the soil, although presumably in lower quantities.

Testing the viability of dry-stored seeds from collections offers an excellent opportunity to identify ‘false transient’ species. *Ex situ* dry storage conditions alter natural aging processes to some extent (Milberg, 1994), yet, seed longevity under dry storage and rapid aging tests significantly positively correlates with soil seed persistence (Bekker *et al*., 2003; Long *et al*., 2015). This correlation implies an intrinsic, genetic basis for persistence through resistance to aging (Bekker *et al*., 2003; Long *et al*., 2008; Probert *et al*., 2009). Based on all these, ‘false transient’ seeds may have low detection probability in the soil caused by methodological issues, low abundance or environmental filters, but they should survive a long-term *ex situ* storage. In contrast, ‘true transient’ seeds are physiologically not prepared to survive a longer period of dry storage.

Building on the demonstrated link between *ex situ* seed longevity and SSB persistence (Bekker *et al*., 2003; Long *et al*., 2015), *ex situ* seed longevity data can also be used to reassess assumptions regarding the relationship between seed persistence and seed size or seed shape. While numerous studies report a negative correlation between seed size and SSB persistence (e.g., Thompson *et al*., 1993; Wang *et al*., 2024; Törő-Szijgyártó *et al*., 2026), we must also consider that low seed production decreases the probability of detecting a species in the SSB (Metzner *et al*., 2017; Saatkamp *et al*., 2009). Therefore, because of the trade-off between seed size and seed number (Leishman, 2001; Moles and Westoby, 2006), the apparent transience of larger seeds in SSB surveys might be an artefact resulting from their lower initial density and, consequently, their lower detection probability (Saatkamp *et al*., 2009). This assumption is supported by laboratory experiments showing no correlation between seed mass and longevity either under gene bank conditions (Pritchard and Dickie, 2003, Walters *et al*., 2005) or in rapid aging tests (Probert *et al*., 2009).

As a first test of the ‘false transience’ hypothesis introduced here, we conducted a germination experiment using dry-stored seeds from a seed collection. Focusing exclusively on plant species considered to have transient seeds based on former field SSB studies, we aimed to answer two main questions: (i) Do species classified as transient include a subset of species capable of retaining germinability after more than 10 years of *ex situ* dry storage (‘false transients’)? (ii) If so, how do the seed mass and seed production of such ‘false transient’ species compare to those of non-germinating transient species (presumably ‘true transients’)?

## Materials & Methods

The seeds were sourced from the seed collection of the HUN-REN-UD Functional and Restoration Ecology Research Group at the University of Debrecen (Hungary). Seeds in this collection are dry-stored under room conditions, with temperature and humidity not closely regulated at the storage location. The seeds experienced a relatively broad range of temperature (approximately 15°C–30°C) and relative humidity. The collection itself contains 1469 species, of which a subset of species were selected for the present experiment. The criteria for including a species in the germination tests were as follows: (i) More than 10 years old seeds of the species were available in the seed collection in sufficient numbers. The mean age of the tested seeds was 15.7 years, ranging from 10 years to 29 years. (ii) The species’ seeds are considered to be transient based on field SSB studies. We extracted data on seed bank persistence primarily from the LEDA Traitbase (Kleyer *et al*., 2008). For species lacking seed bank persistence data in the LEDA Traitbase, we used data from the Pannonian Database of Plant Traits (PADAPT, Sonkoly *et al*., 2023). Then, similar to the seed longevity index of Bekker *et al*. (1998), we calculated the seed bank persistence index (SBPI) to express the proportion of studies indicating persistence for each species. SBPI ranges from 0 to 1, zero meaning that all available studies indicate a transient seed bank for the species, while 1 means that all the studies indicate a persistent seed bank. Species with SBPI<0.2 were considered transient, thus selected for the present study (note that 52% of the studied species had an SBPI=0).

Based on these criteria, we selected 103 species for the germination tests (Supplementary Table S1). Twenty intact seeds of each species were placed on moist filter paper in Petri dishes and incubated under controlled conditions in a climate chamber (Aralab FitoClima 600). Experimental conditions were identical for all species and across all germination trials, but we applied different dormancy-breaking treatments in each trial (see Table S1) in an attempt to minimise the number of species not germinating despite being viable. The seeds were kept under an alternating temperature and light regime consisting of 12 hours of darkness at 15°C followed by 12 hours of light at 25°C, at 40% relative humidity. The germination period lasted 7 weeks in each trial. Germinated seeds were counted, recorded, and removed weekly. Water was replenished regularly as needed.

Three separate trials were conducted (see Table S1) because it was not possible to test all species at once due to the limited capacity of the climate chamber. For the first germination trial, we selected 70 species that could reasonably be assumed to germinate without stratification based on their dormancy type (Baskin and Baskin, 2000) or literature data about their germination requirements. In the case of physiologically dormant (PD) species, long-term dry storage can be expected to break dormancy (i.e., after-ripening, Baskin and Baskin, 2014). Therefore, PD species were included in the first germination trial without any prior dormancy-breaking treatment, unless literature records specifically indicated a need for pre-germination treatment for the species (see the ‘Literature used for determining pre-germination treatment’ column in Table S1). Species with morphophysiological dormancy (MPD) were included in the first germination trial only if literature results indicated that the species germinates without treatment (*Caltha palustris* for example). Furthermore, physically dormant (PY) species were also included already in the first trial, because mechanical scarification provides an easy and rapid method for breaking dormancy in such hard-seeded species. Species that needed stratification, chemical scarification or seed coat removal according to literature records or based on their dormancy type were included in the second and third trial (see Table S1).

In the second germination trial, we included species that were considered to need a dormancy-breaking treatment other than mechanical scarification. Furthermore, we also included species, which were already included in the first trial, in case they did not germinate without a dormancy-breaking treatment and a sufficient number of seeds was still available for testing. Altogether 60 species were tested in the second trial, some of which were also included in the first one. In this trial, most species received wet cold stratification, while some received seed coat removal or chemical scarification using bleach, based on literature results (see Table S1). Finally, we conducted a third germination trial with 51 species, which were all included in the second trial as well, but in this case, they received dry cold stratification instead of wet cold stratification.

Mechanical scarification was carried out by gently rubbing the seeds with sandpaper. Wet cold stratification was carried out by putting the seeds in zip-lock bags alongside a wet paper towel, and placing the bags at 4°C in the fridge for 3 months. Dry cold stratification also meant 3 months at 4°C in the fridge, but under dry conditions in Eppendorf tubes. Chemical scarification was carried out by soaking the seeds in 2% bleach (NaHClO) solution for two days (Rosbakh *et al*., 2019).

Thousand-seed mass data for the tested species were obtained from PADAPT (Sonkoly *et al*., 2023). We note that the majority of the thousand-seed mass data in PADAPT were measured on the same seed lots that were subsequently used in the present germination experiments. To characterise the seed production of the species, we obtained seed number data from the LEDA Traitbase (Kleyer *et al*., 2008) on 10 August 2026. The dataset was restricted to seed number measurements ‘per ramet/tussock or individual plant’, excluding measurements per inflorescence, stem or square metre. Due to the low number of germinating species, we did not perform formal statistical analyses to compare the seed mass and seed production of ‘false transient’ and non-germinating transient species. Instead, we visualised the seed mass and seed number range of all tested species, highlighting the positions of the ’false transient’ species within these distributions. Plots were generated in R version 4.3.2 (R Core Team, 2023). Nomenclature follows Plants of the World Online (POWO, 2026).

## Results

From the 103 tested species, eight species (7.8%) germinated successfully: *Cynosurus cristatus*, *Holcus mollis*, *Rabelera holostea*, *Trifolium medium*, *Vicia cracca*, *V. sativa* subsp. *nigra* (syn. *V. angustifolia*), *V. sepium*, and *Vincetoxicum hirundinaria* (Table 1), all in the first germination trial. Four of the eight species that successfully germinated did so without a dormancy-breaking treatment, while the other four species received mechanical scarification (Fabaceae species). Neither wet nor dry cold stratification, chemical scarification or seed coat removal induced germination in any of the species.

**Table 1.** Information on the species whose seeds germinated in our experiment. Notations: SBPI – seed bank persistence index calculated based on seed bank persistence data from the LEDA Traitbase (Kleyer *et al*., 2008). *SBPI based on data from PADAPT (Sonkoly *et al*., 2023). TSM – thousand-seed mass data from the Pannonian Database of Plant Traits (PADAPT, Sonkoly *et al*., 2023). Seed number – Mean of seed number data for the species from the LEDA Traitbase restricted to measurements ‘per ramet/tussock or individual plant’.

| Species | Family | Seed<br>age<br>(year) | Germination<br>(%) | SBPI | TSM<br>(g) | Seed<br>number | Applied<br>treatment |
| --- | --- | --- | --- | --- | --- | --- | --- |
| *Cynosurus*<br>*cristatus* | Poaceae | 14 | 30 | 0.011 | 0.52 | NA | none |
| *Holcus mollis* | Poaceae | 14 | 10 | 0.033 | 0.299 | 424 | none |
| *Rabelera*<br>*holostea* | Caryophyllaceae | 14 | 25 | 0.095 | 3.034 | 19 | none |
| *Trifolium*<br>*medium* | Fabaceae | 11 | 90 | 0 | 1.842 | 70 | mechanical<br>scarification |
| *Vicia cracca* | Fabaceae | 19 | 40 | 0.043 | 12.759 | 154 | mechanical<br>scarification |
| *Vicia sativa*<br>*subsp. nigra* | Fabaceae | 16 | 15 | 0* | 11.094 | 175 | mechanical<br>scarification |
| *Vicia sepium* | Fabaceae | 16 | 50 | 0.032 | 19.661 | 77 | mechanical<br>scarification |
| *Vincetoxicum*<br>*hirundinaria* | Asclepiadaceae | 14 | 5 | 0 | 7.055 | NA | none |

The majority of species with successful seed germination after 11-19 years of dry storage had relatively large seeds: the seed mass of 75% of the germinating species was above the median seed mass of the tested species, while 50% of the germinating species had seed masses in the upper quartile (Fig. 1). Although seed number data was not available for two of the germinating species, all the other germinating species had seed numbers below the median seed number of the tested species (Fig. 1).

**Figure 1.**
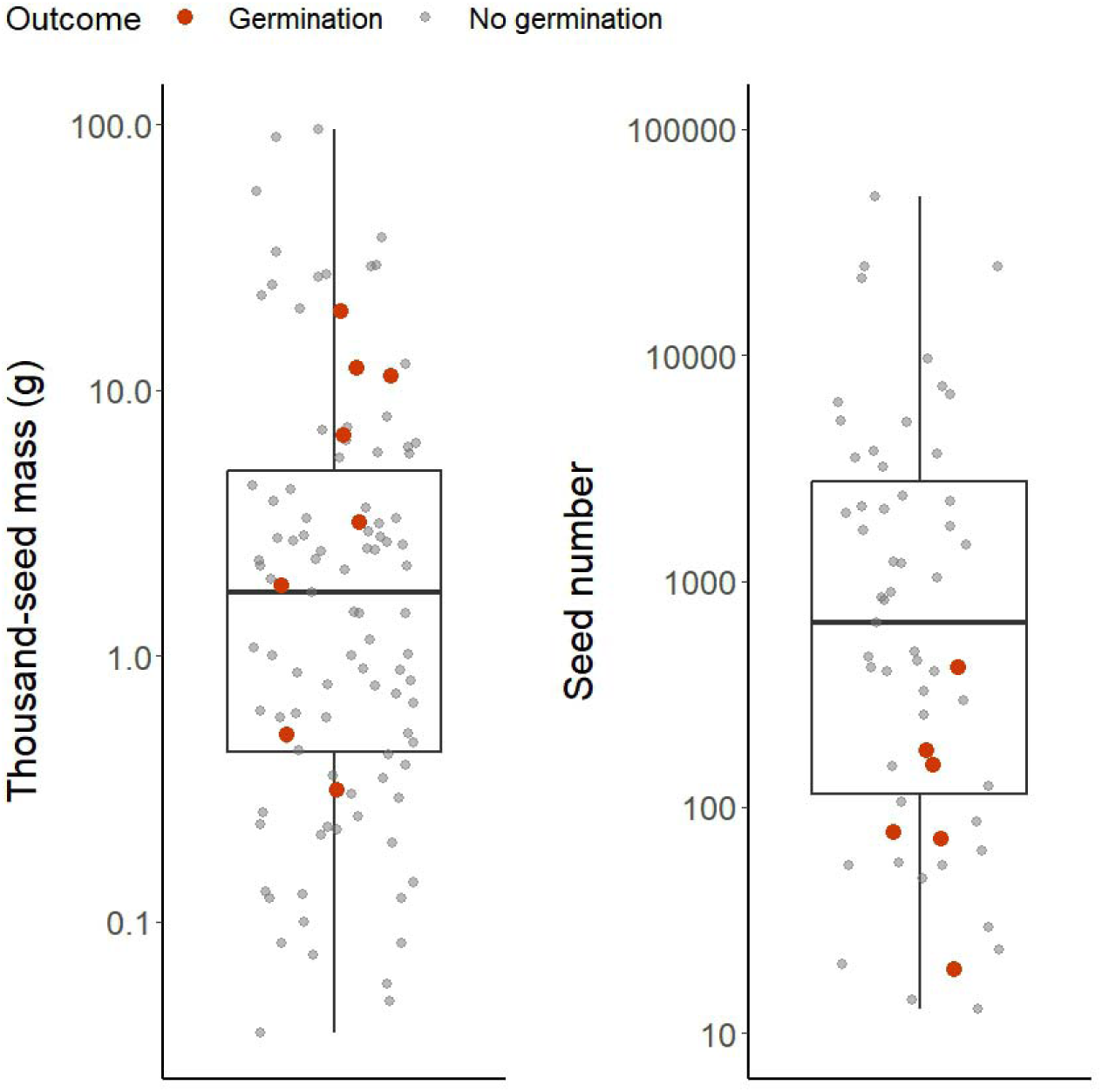
The thousand-seed mass and seed number of species that germinated (‘false transients’) compared to the full range of seed mass and seed number of the tested species.

## Discussion

Here we introduced the ’false transience’ hypothesis, which proposes that a subset of species classified as transient by standard field SSB surveys actually possess the capacity for long-term seed survival. The results of our germination test employing seeds dry-stored for 10-29 years serve as partial support for this hypothesis. Almost 10% of the transient species included in our study successfully germinated after more than 10 years of *ex situ* dry storage, suggesting that a considerable share of species classified as transient may actually be persistent. SSB studies are prone to artefacts caused e.g. by low-abundance species or those having low seed production being misclassified as transient (Thompson *et al*., 1997; Saatkamp *et al*., 2009), while our *ex situ* screening method excluded the confounding effect of detection probability. This suggests that the perceived transience of these species may be caused by sampling and methodological limitations, alongside environmental filters, rather than physiological constraints of the seed to preserve its germination potential for a longer period (‘false transience’). Species that did not show germinability in our experimental tests are more likely to be true transients, however, as dormancy-breaking treatments or germination conditions may have been suboptimal for some species, we cannot draw exact conclusions about the seed longevity of these species.

Environmental conditions experienced by the seeds can have a strong impact on their longevity (Abedi *et al*., 2014; Chen *et al*., 2021), and *ex situ* storage conditions may substantially differ from those experienced by the seeds in the field. Dry-stored seeds may have an advantage compared to seeds kept under field (natural) conditions, since dry storage likely reduces microbial activity and the exposure to seed pathogens they would encounter in the soil (Leishman *et al*., 2000; Schafer and Kotanen, 2003). On the other hand, seeds tested by us experienced a relatively wide range of temperatures (approximately 15°C–30°C) during storage, representing a relatively warm environment that can speed up biochemical processes associated with seed aging, such as enzyme degradation and free radical production (Walters, 1998). Therefore, although the seeds were not exposed to all factors contributing to viability loss (Chambers and MacMahon, 1994), the conditions they experienced are suboptimal for preserving seed viability (Albani Rocchetti *et al*., 2022a). This suggests that the seeds of species successfully germinated in our experiment after more than 10 years of dry storage have an innate ability to remain viable for an extended period of time, even under suboptimal conditions.

Although there is no correlation between seed persistence in the soil under field conditions and seed longevity under dry cold storage conditions (15% RH, -18 to -20°C) in gene banks, such as the Millenium Seed Bank (Walters *et al*., 2005), the highly controlled and very cold conditions in artificial seed banks are far from the conditions experienced by seeds in the field. Conditions during controlled aging tests (rapid aging), i.e., relatively high temperatures and relative humidity levels, are much closer to what seeds actually experience on or near the soil surface under field conditions. Controlled aging conditions test the ability of seeds to withstand moisture and temperature stress, reflecting the same biochemical mechanisms that influence seed persistence under field conditions (Long *et al*., 2008). In accordance with this, seed longevity in controlled aging tests is positively correlated with seed persistence in the soil (Long *et al*., 2008). As the storage conditions of the seeds in our study are closer to controlled aging tests than to artificial seed bank (gene bank) conditions, the longevity of seeds in our study likely also reflects seed persistence in the soil.

Four out of the five tested Fabaceae species showed seed germinability after extended dry storage, with the exception of *Lathyrus pratensis*. Therefore, as expected, our results confirm ‘false transience’ for most of the Fabaceae species tested in our study. Our finding is consistent with the observation that many Fabaceae species have particularly long-term seed viability (e.g., Leino and Edqvist, 2010; Molnár V. *et al*., 2015), in some cases persisting for extremely long periods (Daws *et al*., 2007). This extended seed longevity is mainly driven by their impermeable seed coat, which protects the embryo from humidity fluctuations, resulting in physical dormancy (Baskin and Baskin, 2014; Hay *et al*., 2023). The lack of germination in *L. pratensis* despite mechanical scarification is in sharp contrast to the successful germination of other Fabaceae species. It is of course possible that *L. pratensis* is a ‘true transient’ species despite having a hard seed coat. On the other hand, the apparent mortality of its seeds under dry storage may also be explained by the fact that hard seed coats require full maturation to effectively protect the seeds from humidity fluctuations, therefore seeds collected too early frequently exhibit poor storage potential (Hay and Probert, 1995; Ali *et al*., 2007). Furthermore, the relatively low number of seeds tested per species may also lead to overlooking viability in some seed lots.

However, in our experiment, even some species without hard, protective seed coats and physical dormancy (*Cynosurus cristatus*, *Holcus mollis*, *Rabelera holostea*, *Vincetoxicum hirundinaria*) showed germinability after 14 years of dry storage. For example, the successful germination of *V. hirundinaria* after 14 years of dry storage is in agreement with the finding that 18-years-old seeds of the congeneric *V. pycnotelma* from herbarium also showed considerable germination (Nakahama, 2015). As all non-Fabaceae species that successfully germinated in our experiment did so without a dormancy-breaking treatment, after-ripening in the form of long-term dry storage seems to be enough to break the dormancy of their seeds. This suggests that these species likely have non-deep physiological dormancy (Baskin and Baskin, 2014). However, their germination rates remained significantly lower (17.5% on average) than that of the physically dormant Fabaceae species (49% on average, up to 90%). Albani Rocchetti *et al*. (2024) also found that germination of old seeds from herbarium specimens sometimes occurs in plant families that do not have physical dormancy. In their study, old herbarium seeds belonging to 12 different families showed germination, indicating that extended seed longevity is possible in several plant families, and not just in families characterised by physical dormancy. In accordance with our findings, they also mentioned that some of the species that germinated successfully in their experiment were thought to have medium or short-lived seeds (Albani Rocchetti *et al*., 2022b, 2024).

On the other hand, our experiment did not disprove transient classification for most species without a hard seed coat and physical dormancy. Despite using targeted dormancy-breaking treatments (cold stratification, chemical scarification, etc.) to reduce the chances of missing viable seeds, germination was zero in most but not all species without physical dormancy. Although our findings increase the likeliness that these species are ‘true transients’, we cannot draw definite conclusions about the persistent versus transient nature of the species not germinating in our experiment, as the germination conditions may have been suboptimal for some of them and the relatively low total number of seeds tested per species (20–60 seeds depending on the species) may also lead to missing potential long-term seed viability in some of them. Furthermore, as more than half of the tested seeds were 15 years old or even older (the mean age of seeds was 15.7 years), the lack of germination does not necessarily indicate an inability to remain viable for several years. These species may still be able to persist in the soil for multiple years, although for a shorter period than the age of the tested seeds. Future studies aiming to identify ‘false transient’ species should test germination under a range of light and temperature conditions or ideally use TTC viability tests (França-Neto and Krzyzanowski, 2022), as these approaches would allow more definitive conclusions regarding seed longevity.

We also aimed to assess how the seed mass and seed production of ‘false transient’ species (those that germinated in our study) compare with the seed mass and seed production of those that did not germinate after more than 10 years of dry storage. The position of the germinating species within the full range of seed mass and seed number spectra observed across the tested species provide evidence that ‘false transients’ are frequently found among the large-seeded species and those having low seed production. The prevalence of large-seededness among ‘false transient’, thus persistent species in our study is not in agreement with the reported association between small seed size and SSB persistence (e.g., Thompson *et al*., 1993; Wang *et al*., 2024; Törő-Szijgyártó *et al*., 2026). In addition, ‘false transient’ species tended to have low seed numbers compared to the whole set of studied species, which were previously all classified as transient. These results are consistent with the hypothesis that the commonly reported greater persistence of small-seeded species may largely reflect a detection artefact rather than an intrinsic relationship between seed size and seed longevity (Saatkamp *et al*., 2009). Larger seeds are generally produced in lower numbers than smaller seeds due to the trade-off between seed size and seed production (Leishman, 2001; Moles and Westoby, 2006), and species producing less seeds tend to accumulate lower seed densities in the soil (Bekker *et al*., 2003). As a result, they are less likely to be detected in SSB studies than small-seeded species (Thompson *et al*., 2003), potentially creating the impression that large-seeded species are less persistent than they actually are. In addition, larger seeds move more slowly downward through the soil compared to smaller seeds (Bekker *et al*., 1998), and may therefore be underrepresented in deeper soil layers, which are thought to contain the majority of persistent seeds. Consequently, SSB surveys may underestimate not only the potential longevity of larger-seeded species, but also their realised (ecological) seed persistence, simply because their lower seed densities reduce the probability of their detection, especially in deeper soil layers (Thompson *et al*., 1997). Our findings are in agreement with the results of other experimental studies, which found no correlation between seed size and seed longevity (Pritchard and Dickie, 2003; Walters *et al*., 2005; Probert *et al*., 2009) or seed bank persistence (Saatkamp *et al*., 2011; Zani and Müller, 2017).

Our findings suggest that persistence classifications based solely on field SSB studies cannot conclusively establish the transient nature of seeds, and they definitely should not be used to predict *ex situ* storage potential. Recognising this ‘cryptic seed persistence’ has important implications for conservation management: species categorised as transient in SSB studies are generally considered unable to reestablish from the soil seed bank, and conservation and restoration decisions are also based on this information (see e.g., Richter and Stromberg, 2005; Wang *et al*., 2020). Based on our results, restoration may count on at least some species categorised as transient. Therefore, to prevent misguided decisions and wasted resources, conservation strategies should distinguish between ‘true transients’ and ‘false transients’. Furthermore, to reduce the risk of erroneously categorising species as transient in SSB studies, we suggest establishing a vegetation frequency or cover threshold below which species are not assigned a persistence category if their seeds are not detected in the samples (see a similar approach in Török *et al*., 2009).

## Conclusions

We can conclude that the apparent absence of certain species from the SSB may only reflect ecological filters (such as the seed size/number trade-off) rather than their seeds’ inherent physiological inability to survive. Relying solely on field SSB data to infer seed longevity can be very misleading, as what we observe in the field is a net ecological effect, not a direct measure of potential longevity. Species that germinated in our experiment are most probably ‘false transients’ and their transient classification is most likely an artefact of low detection probability driven mostly by low seed production and/or low abundance in the vegetation. However, for species that did not show germinability in our experiment, it remains unclear whether their observed transience is a physiological reality or a sampling artefact. To resolve this question, targeted, species-specific data on seed longevity will likely be necessary. Ultimately, our findings reinforce that observed seed bank persistence and seed longevity are distinct phenomena and should not be confused (Saatkamp *et al*., 2009). Resolving whether seed traits like seed size and shape genuinely dictate longevity and persistence or merely influence detection probability is vital for understanding seed bank dynamics; therefore, larger-scale studies of this issue would be essential to validate our findings.

## Supporting information

Detailed information about the species included in the germination experiments, pre-germination treatments, and germination outcomes.

## Supplementary Information

**Table S1:** Detailed information about the species included in the germination experiments, pre-germination treatments, and germination outcomes.

## Author Contributions (CRediT)

J.S.: conceptualization, methodology, data curation, formal analysis, writing – original draft. V.T.-S.: methodology, investigation, writing – original draft. E.R.: conceptualization, writing – review and editing. K.T.: investigation. G.K.-V.: investigation. S.M.: investigation. E.T.-S.: investigation. P.T.: conceptualization, methodology, funding acquisition, writing – review and editing.

## Data Availability

All the data generated for and used in this study are available in Supplementary Table S1.

## Acknowledgements

We are grateful for colleagues who provided seeds for the seed collection used in this study.

## AI Assistance Acknowledgements

We acknowledge the use of Google Gemini (version Gemini 3.1 Pro) for refining the language and grammar of specific sections of this manuscript. The authors reviewed, edited, and take full responsibility for the final content.

